# NHR-23 and SPE-44 regulate distinct sets of genes during *C. elegans* spermatogenesis

**DOI:** 10.1101/2022.06.24.497528

**Authors:** James Matthew Ragle, Kayleigh N. Morrison, An A. Vo, Zoe E. Johnson, Javier Hernandez Lopez, Andreas Rechtsteiner, Diane C. Shakes, Jordan D. Ward

## Abstract

Spermatogenesis is the process through which mature male gametes are formed and is necessary for transmission of genetic information. While much work has established how sperm fate is promoted and maintained, less is known about how the sperm morphogenesis program is executed. We previously identified a novel role for the nuclear hormone receptor transcription factor, NHR-23, in promoting *C. elegans* spermatogenesis. Depletion of NHR-23 along with SPE-44, another transcription factor that promotes spermatogenesis, caused additive phenotypes. Through RNA-seq, we determined that NHR-23 and SPE-44 regulate distinct sets of genes. Depletion of both NHR-23 and SPE-44 produced yet another set of differentially regulated genes. NHR-23- regulated genes are enriched in phosphatases, consistent with the switch in spermatids to post-translational regulation following genome quiescence. In the parasitic nematode *Ascaris suum*, MFP1 and MFP2 control the polymerization of Major Sperm Protein, the molecule that drives sperm motility and serves as a signal to promote ovulation. NHR-23 and SPE-44 regulate a number of MFP2 paralogs, and NHR-23 depletion caused defective localization of MSD/MFP1 and NSPH-2/MFP2. Although NHR-23 and SPE-44 do not transcriptionally regulate the casein kinase gene *spe-6,* a key regulator of sperm development, SPE-6 protein is lost following NHR-23+SPE-44 depletion. Together, these experiments provide the first mechanistic insight into how NHR-23 promotes spermatogenesis and an entry point to understanding the synthetic genetic interaction between *nhr-23* and *spe-44*.

## INTRODUCTION

Spermatogenesis involves a cascade of morphogenetic events to produce the highly specialized, haploid, motile gametes essential for sexual reproduction. Universal features of spermatogenesis include the biosynthesis of sperm-specific proteins and assembly of sperm-specific complexes, preparing for and progressing through the meiotic divisions, and a streamlining step during which cells discard cytoplasmic components that are no longer needed. At species-specific points during sperm development, transcription ceases; the final steps of differentiation and mature sperm function occur in the absence of transcription. The sperm’s genome is protected from environmental and metabolic damage through a spermatogenesis-specific process of chromatin remodeling and hypercompaction. These processes are all essential for faithful transmission of the sperm genome to offspring.

*C. elegans* is a powerful model to study spermatogenesis. Within the self-fertile hermaphrodites, uncommitted germ cells have the capacity to undergo either spermatogenesis or oogenesis; but in the normal process, the hermaphroditic germline makes a one-time switch from spermatogenesis to oogenesis as animals enter adulthood. Analysis of the accompanying changes provides insight into how sperm and egg fates are promoted and maintained. Alternatively, analysis of males allows the study of continual spermatogenesis. Both hermaphrodite and male germlines are polarized organs; the uncommitted germ cells proliferate mitotically at the distal end before committing to entering meiotic prophase and commitment to the sperm fate. The events of meiosis occur in a linear fashion along the length of the gonad, resulting in the production of haploid spermatids. These include homologous chromosome pairing, double-stranded breaks in the DNA promoting recombination, and two meiotic divisions (Shakes *et al*. 2009; Chu and Shakes 2013; Ellis and Stanfield 2014; Griswold 2016). During *C. elegans* spermatogenesis, general transcription ceases near the end of meiotic prophase, and there is no burst of post-meiotic transcription as occurs in humans and Drosophila (Bettegowda and Wilkinson 2010; Chu and Shakes 2013). Translation ceases shortly after anaphase II, as not only ribosomes but also tubulin, actin, and endoplasmic reticulum are partitioned away from the budding spermatids and into a structure known as the residual body (Chu and Shakes 2013; Fig. 1). Lacking typical cytoskeletal proteins, nematode sperm motility is driven by the assembly/disassembly dynamics of the 14 kDa nematode-specific Major Sperm Protein (MSP) within the pseudopod of the crawling spermatozoa. The early cessation of both transcription and translation requires all sperm-specific components to be synthesized during meiotic prophase prior to their developmental deployment. During this critical window within developing spermatocytes, newly synthesized MSP assembles into large paracrystalline fibrous bodies (FBs) that form in association with Golgi-derived membranous organelles (MOs; Fig. 1). This form of MSP packaging facilitates the efficient partitioning of MSP to spermatids.

**Fig 1.**
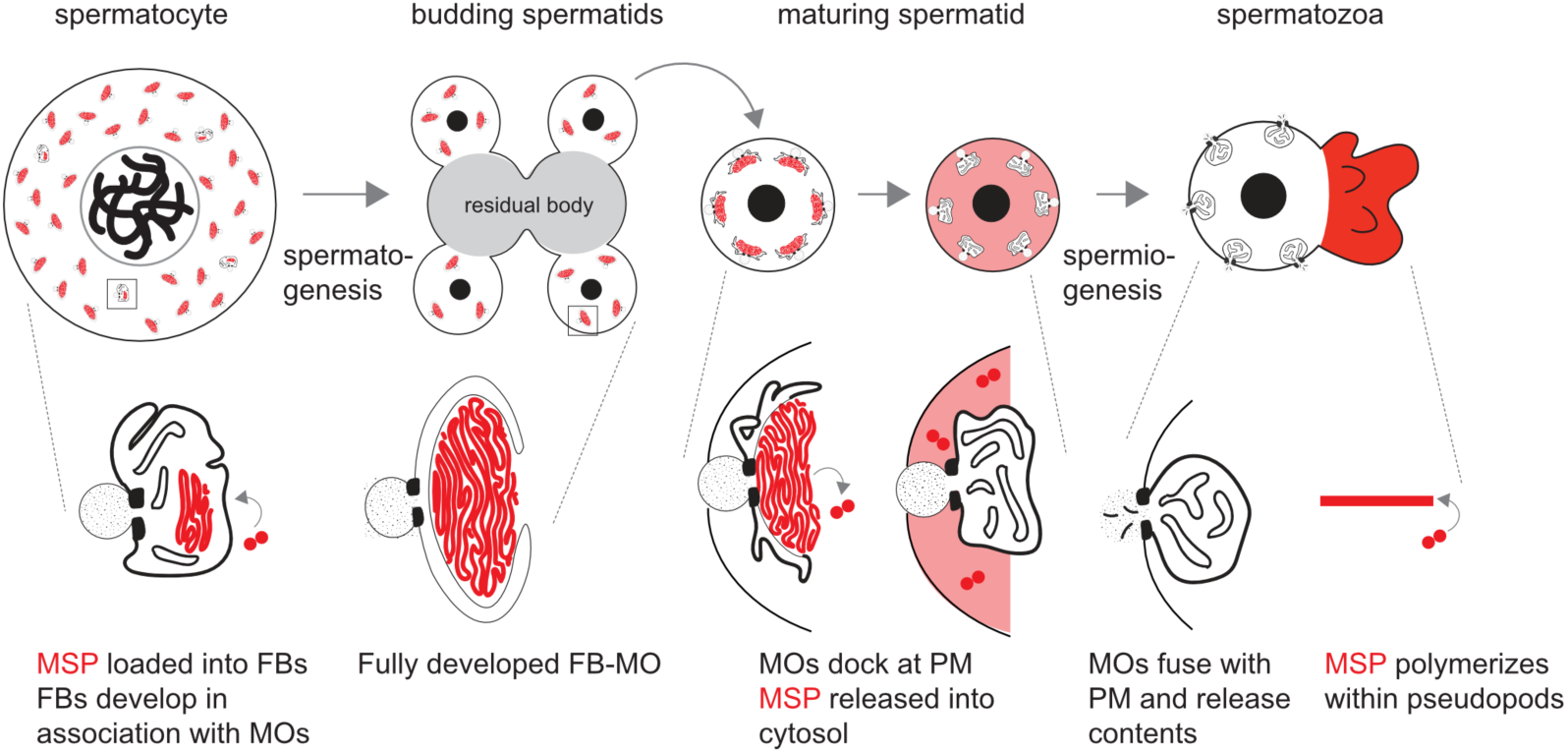
Overview of Major Sperm Protein (MSP) dynamics during spermatogenesis. (Top) Spermatocytes undergo reductive cell division during spermatogenesis. During this process, MSP is packaged and sequestered in FB-MOs. Following anaphase II, components which are no longer needed partition to a central residual body, which is discarded. Spermatids bud off of the residual body, mature, and upon activation, undergo morphological changes during spermiogenesis. The differentiated spermatozoa contain a cell body and pseudopodia. (Bottom) Within spermatocytes, fibrous bodies (FBs) develop on the cytosolic face of the Golgi-derived membranous organelle (MO). During the budding division, FB-MO complexes partition to the spermatids. In maturing spermatids, MOs dock with the plasma membrane as the FBs disassemble and release MSP dimers into the cytosol. During sperm activation, MSP localizes to the pseudopod, where it polymerizes, and MOs fuse with the plasma membrane.

The transcription and translational programs that govern *C. elegans* spermatogenesis must not only drive differentiation of spermatocytes and oocytes, but they must also create the materials that spermatocytes subsequently need for sperm differentiation and sperm function. How early acting gene regulatory networks provision spermatocytes for the extended process of spermatogenesis and sperm morphogenesis remains an open question. To date, two transcription factors have been characterized as critical regulators of *C. elegans* spermatogenesis, SPE-44 and NHR-23. SPE-44 was first identified by its enrichment in sperm-producing germlines. Further investigation revealed that *spe-44* null spermatocytes fail to load MSP into FB-MOs, and they are arrested as terminal spermatocytes during the second meiotic division (Kulkarni *et al*. 2012). *spe-44* regulates a broad set of genes that are enriched in spermatogenic germlines (Kulkarni *et al*. 2012). Consistent with the MSP loading defects, these genes include MSP genes and members of the small sperm-specific protein family, which promote MSP polymerization (Kulkarni *et al*. 2012). *spe-44* also regulates several genes previously identified as being necessary for spermatogenesis through forward genetic screens for spermatogenesis defective (*spe*) and fertilization-defective (*fer*) genes (Kulkarni *et al*. 2012). However, there were numerous *spe* and *fer* genes which were not regulated by *spe-44* (Kulkarni *et al*. 2012). In terms of regulatory hierarchies, *spe-44* was also shown to regulate the transcription factor, *elt-1* (Kulkarni *et al*. 2012), which is a direct regulator of MSP gene expression (del Castillo-Olivares *et al*. 2009).

We recently discovered that the nuclear hormone receptor NHR-23, a critical regulator of nematode molting, is also expressed in primary spermatocytes and necessary for spermatogenesis (Ragle *et al*. 2020). NHR-23 and SPE-44 expression patterns almost completely overlap (Ragle et al. 2020). Both proteins label the autosomes but not the X-chromosome from the beginning of meiotic prophase until global transcription ceases and spermatocytes enter the karyosome stage. Germline-specific NHR-23 depletion causes a terminal spermatocyte arrest during the first meiotic division (Ragle et al. 2020). Both *spe-44* and nhr-23 spermatocytes fail to load MSP into FBs, however, the Golgi-derived MOs of the FB-MO complex still assemble in the absence of NHR-23 (Ragle et al. 2020). We determined that SPE-44 and NHR-23 are in separate genetic pathways because they do not regulate one another, and because NHR-23 and SPE-44 double depletion produce more severe synthetic phenotypes such as a nearly complete loss of MO differentiation markers and severe germline defects (Ragle *et al*. 2020). Exactly how NHR-23 promotes spermatogenesis was not clear. Here, we use a combination of transcriptomics, genome editing, and cytology to address this question and gain insight into the synthetic genetic interaction between *spe-44* and *nhr-23*.

## MATERIALS AND METHODS

### Strains and culture

*C. elegans* were cultured as originally described (Brenner 1974), except that worms were grown on MYOB media instead of NGM. MYOB agar was made as previously described (Church *et al*. 1995). Brood sizes were performed picking L4 larvae to individual wells of a 6-well plate seeded with OP50 and incubating the plate at 20°C. Animals were transferred to new plates daily over 4 days. Two days post-transfer, the number of hatched progeny and unfertilized eggs were scored. We scored unfertilized eggs over the first three days as on the 4th day animals frequently ran out of sperm.

The following strains were used in this study:

| Strain | Genotype | Source/reference |
| --- | --- | --- |
| N2 | Wild type | CGC |
| JDW85 | *wrdSi14[mex-5p::TIR1:F2A:mTagBFP2:tbb-2 3'UTR, l:-5.32]; him-5(e1490) V.* | Ragle *et al.*, 2020 |
| JDW86 | *wrdSi15[mex-5p::TIR1:F2A:mTagBFP2:tbb-2 3'UTR, l:-5.32]; spe-44(fx110[spe-44::AID\*]) IV; him-5(e1490) V.* | Ragle *et al.*, 2020 |
| JDW92 | *wrdSi19[mex-5p::TIR1:F2A:mTagBFP2:tbb-2 3'UTR, l:-5.32], nhr-23(kry61(nhr-23::AID\*-TEV-3xFLAG)) I; him-5(e1490) V.* | Ragle *et al.*, 2020 |
| JDW146 | *wrdSi19[mex-5p::TIR1:F2A:mTagBFP2:tbb-2 3'UTR, l:-5.32], nhr-23(kry61(nhr-23::AID\*-TEV-3xFLAG)) I; spe-44(fx110[spe-44::degron]) IV ; him-5(e1490) V.* | Ragle *et al.*, 2020 |
| JDW307 | *nsph-3.2(wrd56[C04G2.9::exon 1 STOP]) IV* | This study |
| JDW308 | *nsph-2(wrd57[ZK265.3::exon 1 STOP]) I* | This study |
| JDW320 | *nsph-2(wrd57[ZK265.3::exon 1 STOP]) I; nsph-3.2(wrd56[C04G2.9::exon 1 STOP]) IV* | This study |
| JDW339 | *nsph-2(wrd57[ZK265.3::exon 1 STOP]) I; nsph-1.1(wrd67[K08F8.5::exon 2 STOP]) II* | This study |
| JDW358 | *ZK596.2(wrd71[exon 1 STOP]) III* | This study |
| JDW361 | *ttbk-5(wrd72[exon 1 STOP]) IV* | This study |
| JDW367 | *nsph-4.3(wrd74[T04A6.4 exon 1 STOP]) III* | This study |
| JDW374 | *ZK757.2(wrd77[exon-1 STOP]) III* | This study |
| JDW403 | *nsph-4.1(wrd92[whole gene deletion]) IV* | This study |
| JDW404 | *nsph-4.2(wrd93[whole gene deletion]) IV* | This study |
| JDW427 | *nsph-4.1(wrd92[whole gene deletion]), nsph-4.2(wrd97[whole gene deletion]) IV* | This study |
| RV120 | *spe-44(ok1400) dpy-20 (e1282) / let-92 (s677) unc-22 (s7) IV*<br>ok1400 deletion removes 1577 of the 1797 base-pair coding region, including the conserved SAND domain.<br>Presumed null | Kulkarni *et al.*, 2012 |

### RNA-seq and bioinformatics

Adult hermaphrodites of the indicated genotypes were picked onto 6 cm MYOB+4 mM auxin plates and allowed to lay eggs for two hours. The hermaphrodites were removed and the plates were incubated at 20°C for 3 days. 500 L4/young adult males were hand-picked into M9 + 0.05% gelatin, washed 1X and pelleted. Liquid was removed down to 25 ul and 500 µl TRIzol (Invitrogen) was added. The mixture incubated at room temperature for 5 minutes with periodic vortexing. Carcasses were pelleted by centrifugation and supernatant was removed to a clean tube. 100 µl chloroform was added to the supernatant, vortexed for 30 seconds and phases separated by centrifugation at 15,000 RPM for 5 minutes. 250 µl of the upper aqueous phase was removed to a new tube and 350 µl chloroform was added, vortexed and separated as in the previous steps. 250 µl of isopropanol was added to the aqueous phase along with 1 µl Glycoblue (Invitrogen) and vortexed. The solution was centrifuged for 20 minutes at 15,000 RPM and the supernatant was decanted. The pellet was washed with 1 ml 75% ethanol twice and dried at room temperature for 10 minutes. The dried pellet was resuspended in 16 µl nuclease-free water. Concentration was determined with the NanoDrop.

Using the NEXTFLEX® Rapid Directional RNA-Seq Kit 2.0 (BioO/PerkinElmer), polyA-selected RNA was isolated from 300 ng of total RNA according to manufacturer’s instructions. Sequencing libraries were prepared using the same kit according to manufacturer’s instructions. Concentrations were measured on a Qubit Fluorometer (Invitrogen) and fragment size determined on a Bioanalyzer 2100 (Agilent). Libraries were sequenced at the University of California Berkeley QB3 NGS facility on a Nova-seq platform. Raw sequences were mapped to transcriptome version WS220 using Tophat2 (Kim *et al*. 2013). Only reads with one unique mapping were retained, otherwise default options for paired-end mapping were used. HTSeq was used to obtain read counts per transcript (Anders *et al*. 2015). All libraries were sequenced twice and for each transcript read counts from both sequencing runs were added together. DESeq2 was used for read count normalization and differential expression analysis (Love *et al*. 2014). DESeq2’s regularized log transformation with option blind=TRUE was used for read count normalization of protein coding genes prior to principal component analysis (PCA). PCA was performed using the prcomp function in the stats package of R (R Core Team). A Benjamini–Hochberg multiple hypothesis-corrected p-value cutoff of 0.05 was used as a cutoff for significant differential expression. Table S1 contains the processed data with gene name and location, mean expression, log2 fold-change compared to control, and p-value. Protein and cDNA alignments (Fig. S2 and S3) were produced with Clustal Omega (Goujon *et al*. 2010; Sievers *et al*. 2011). Protein alignments were visualized with JalView2 (Waterhouse *et al*. 2009). Venn diagrams were generated using GeneVenn (http://genevenn.sourceforge.net/) and figures were generated in Affinity Designer.

### CRISPR/Cas9-mediated genome editing

Mutations in candidate NHR-23 target genes were generated using homologous recombination of CRISPR/Cas9-induced double-strand breaks (DSBs). We made Cas9 ribonucleoprotein (RNP) complexes (750 ng/µl house-purified Cas9, 50 ng/µl crRNA and 100 ng/µl tracrRNA) by incubating at 37C for 15 minutes. RNPs were mixed with oligonucleotide repair templates (110 ng/µl; Ghanta and Mello 2020; Paix *et al*. 2015; Dokshin *et al*. 2018) and a co-injection marker (either 50 ng/µl pRF4, 1-10 ng/µl pKA411 or 25 ng/µl pSEM229 (El Mouridi *et al*. 2020 p. 1)) to facilitate isolation of successfully injected progeny. Where possible, we selected “GGNGG” crRNA targets as these have been the most robust in our hand and support efficient editing (Farboud and Meyer 2015). All STOP-IN cassettes were designed to be inserted directly at the DNA DSB site. Sequences and descriptions of the crRNAs and repair oligos are provided in Table S2 and S3, respectively. We picked L4 animals, aged them one day at 20°C and injected 6-18 animals for each edit. Injected animals were grown at either 20°C or 25°C until progeny were produced. These progeny were isolated to individual plates to lay eggs. We made genomic lysates from parent animals and genotyped them by polymerase chain reaction using primers that anneal to sites flanking the mutations. As our version of the STOP-IN cassette contains a *Bam*HI site, insertions were identified by diagnostic *BamHI* digestion. Whole gene deletions were identified by PCR product size. Mutations and homozygosity were verified by Sanger sequencing. Genotyping primers are provided in Table S4. Knock-in sequences are provided in File S1.

### Immunocytochemistry

Embryos were isolated from adult hermaphrodites of the indicated genotypes by standard bleaching methods (dx.doi.org/10.17504/protocols.io.j8nlkkyxdl5r/v1) and allowed to hatch overnight in the absence of food. L1 larvae were transferred to multiple OP50 plates for 6-8 hours. The treatment groups were washed off the plates and transferred to MYOB+4 mM auxin plates. Sibling controls were kept on non-auxin plates. Both groups were raised to adulthood at room temperature. Sperm spreads were obtained by dissecting 10-12 male worms in 7 microliters of egg buffer <u>(Edgar 1995)</u> on ColorFrost Plus slides (Fisher Scientific, 12-550) that were additionally coated with poly-L-lysine (Sigma Aldrich, P8290). Worms of different genotypes were dissected on the same slide to ensure that antibody binding conditions were competitive and identical. During analysis, mutants were distinguished by their DAPI-stained nuclear patterns. Separate immunocytology preps were performed with worms of different genotypes on different slides to help identify the phenotypes observed on mixed slides. Samples were freeze-cracked in liquid nitrogen and fixed overnight in −20°C methanol. Specimen preparation and antibody labeling followed established protocols <u>(Shakes *et al*. 2009)</u>. Primary antibodies included: 1:300 rabbit anti-MSD-1 (F44D1.3) polyclonal <u>(Kosinski *et al*. 2005)</u>, 1:200 rabbit anti-MFP2/NSPH-2 (ZK265.3) polyclonal <u>(Morrison *et al*. 2021)</u>, and 1:250 rabbit anti-SPE-6 (Peterson et al., 2021). 1:400 Alexa Fluor Plus 555 goat anti-rabbit IgG (Invitrogen, A32732) was used as the secondary antibody. Final slides were mounted with Fluoro Gel with DABCO (Electron Microscopy Sciences #17985-02) containing DAPI. Images were acquired with epifluorescence using an Olympus BX60 microscope equipped with a QImaging EXi Aqua CCD camera. Photos were taken, merged, and exported for analysis using the program iVision. Image exposures were optimized for control samples and then all samples were imaged with the same exposures and no further image processing so that images would reflect relative intensities.

### Data availability

Strains and plasmids are available upon request. File S1 contains the genomic sequence of wild type and genome edited loci. The RNA-seq data has been deposited in the National Center for Biotechnology Information Gene Expression Omnibus under accession number GSE206676. Table S1 contains the processed data with gene name and location, mean expression, log2 fold-change compared to control, and p-value. Sequences and descriptions of the crRNAs and repair oligos are provided in Table S2 and S3, respectively. Genotyping primers are provided in Table S4. All supporting files will be deposited in Figshare.

## RESULTS

### NHR-23 and SPE-44 regulate distinct sets of target genes

Given that NHR-23 is a transcription factor, we hypothesized that loss of NHR-23- regulated gene expression was mediating the phenotypes we observed when NHR-23 was depleted. We therefore depleted NHR-23, SPE-44, and NHR-23+SPE-44 in male germlines using a *mex-5p::TIR1* transgene that we developed for an auxin-inducible degron toolkit (Ashley *et al*. 2021). We performed bulk RNA-seq on whole male animals to identify differentially regulated genes relative to a *mex-5p::TIR1* control strain grown on auxin. With our sequencing data, we performed principal component analysis of the replicates of all conditions. The first 3 principal components captured 63% of the variance and separated the various conditions. (Fig. 2A,B). Using a p<0.05 cutoff, there were only 70 genes that significantly changed in expression when NHR-23 was depleted (Fig. 2C, Table S5). NHR-23 appeared to function primarily to activate gene expression as 55/70 genes were down-regulated following NHR-23-depletion (Fig. 2C). SPE-44-depletion resulted in 811 differentially expressed genes, with 523 genes being down-regulated (Fig. 2D, Table S6). For all three datasets, we set the cutoff for differentially regulated genes as a >1.5 fold change in expression relative to the control. NHR-23+SPE-44 double depletion produced 554 differentially expressed genes, with 494 being down-regulated (Fig. 2E, Table S7). When compared to an RNA-seq study of spermatogenesis versus oogenesis enriched genes (Ortiz *et al*. 2014), NHR-23-regulated and NHR-23+SPE-44- regulated genes had a high percentage (72.9% (51/70) and 84.3% (467/554), respectively) of sperm-enriched genes ((Fig 2G, H). In contrast, only 37.7% (306/511) of SPE-44-regulated genes were spermatogenesis-enriched (Fig. 2I). A *spe-44* mutant microarray experiment found 50.9% (358/703) genes were spermatogenesis-enriched, though this percentage went to 67.8% (343/506) if only the down-regulated genes were analyzed (Kulkarni *et al*. 2012). When we performed a similar analysis on our SPE-44- downregulated genes we found 53.5% (296/553) were spermatogenesis-enriched (Fig S1A).

**Fig. 2.**
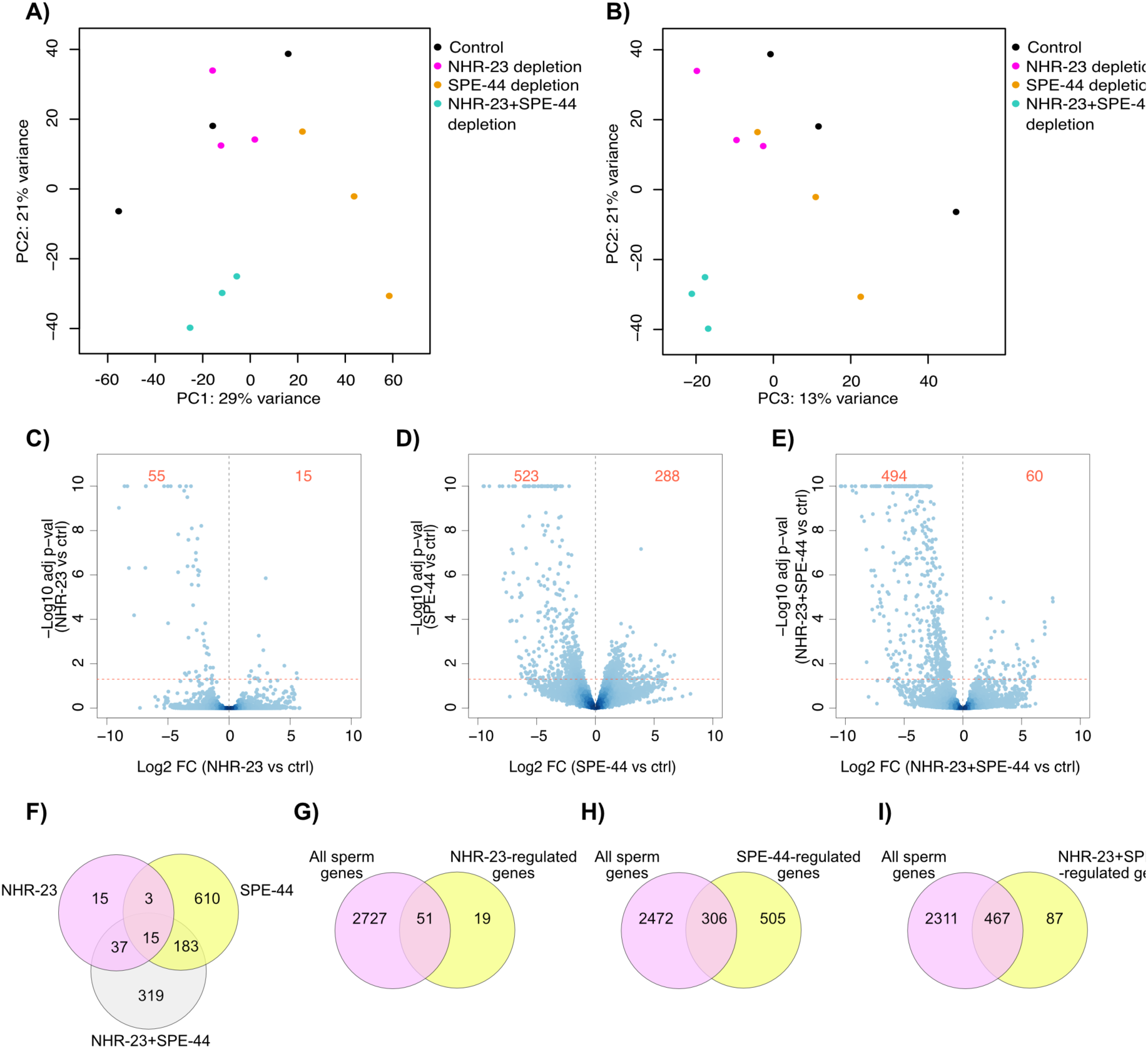
NHR-23 and SPE-44 regulate distinct sets of genes. (A and B) Principal component analysis of RNA-seq data. Legend indicates which colored dots belong to control, NHR-23-depleted, SPE-44- depleted, or NHR-23+SPE-44-depleted biological replicates. (C-E) Volcano plots of the indicated depletion datasets showing Log10 adjusted p-value against Log2 fold-change (FC). The horizontal red dotted line represents our p-value cutoff of 0.05 and the vertical black dotted line indicates a fold-change of 0. The red numbers in each figure indicate the number of differentially down-regulated (left side) and up-regulated (right side) genes in each comparison with a fold-change cutoff >1.5 (log2=0.58) and p<0.05. (F) Venn diagram depicting overlap of differentially regulated genes following NHR-23-, SPE-44, and NHR-23+SPE-44 depletion. Venn diagrams depicting overlap between sperm-enriched genes <u>(Ortiz *et al*. 2014)</u> and differentially-regulated genes following NHR-23 depletion (G), SPE-44-depletion (H), and NHR-23+SPE-44 depletion (I).

We were surprised to find so few differentially-regulated genes following NHR-23 depletion in the germline as *nhr-23* regulates 265 genes in the soma (Kouns *et al*. 2011). NHR-23 regulated genes included a few genes previously implicated in spermatogenesis. NHR-23 regulated the Major Sperm Protein Domain containing proteins MSD-1, MSD-2, and MSD-3 which are poorly characterized paralogs of an MSP polymerization inhibitor in Ascaris, MSP fiber protein 1 (MFP1) (Buttery *et al*. 2003; Grant *et al*. 2005; Kosinski *et al*. 2005; Morrison *et al*. 2021)(Table 1). *smz-2* was also NHR-23 regulated (Table 1). *smz-2* has a paralog (*smz-1)*, and the proteins SMZ-1/2 localize to chromatin during meiosis (Chu et al. 2006). Additionally, RNAi targeting either SMZ gene causes a metaphase arrest during meiosis (Chu *et al*. 2006). In contrast to the short list of NHR-23 regulated genes, we found 811 genes that were differentially expressed upon SPE-44 depletion (Figure 2, Table S6) and 554 genes differentially expressed when both NHR-23 and SPE-44 were depleted (Figure 2, Table S7). Surprisingly, our SPE-44-regulated gene set had limited overlap with a set of *spe-44-*regulated genes identified through a microarray experiment using *spe-44* null L4 males (Kulkarni *et al*. 2012). Only 180 of the 811 SPE-44-regulated genes from our RNA-seq dataset were common to the *spe-44* null microarray dataset (Figure S1B). The transcription factor *elt-1*, a downstream target of *spe-44,* was found in both the microarray and RNA-seq datasets, as was a target validated by promoter reporters in Kulkarni et al. (2012)(F32A11.3 (*spe-18*), referred to as *spe-7* in that paper)(Table 1). Two *msp* genes (*msp-49, msp-63)* were common to the microarray and RNA-seq datasets (Table 1). There were five *spe* genes that were down-regulated in our RNA-seq data, but not in the *spe-44* microarray (Table 1). Three genes encode membrane proteins required for fertilization (*spe-9, spe-42, spe-49*), *spe-43* is required for sperm activation, and *spe-45* is required for sperm to fuse with the oocyte plasma membrane (Singson *et al*. 1998; Zannoni *et al*. 2003; Kroft *et al*. 2005; Nishimura *et al*. 2015; Krauchunas *et al*. 2018; Wilson *et al*. 2018; Takayama *et al*. 2021). While our current focus in this study is on how NHR-23 promotes spermatogenesis, exploring the commonalities and differences between our RNA-seq data and the microarray data is an important future direction to further elucidate how SPE-44 promotes sperm development.

**Table 1.** Differential regulation of select genes implicated in spermatogenesis.

| Gene | Log2 fold change following depletion of: |  |  | Function |
| --- | --- | --- | --- | --- |
|  | NHR-23 | SPE-44 | NHR-23+SPE-44 |  |
| *msd-1* | -2.51<br>(1.02E-06) | - | -3.13<br>(1.62E-11) | MFP1 paralog. Potential role in slowing MSP polymerization |
| *msd-2* | -2.42<br>(6.87E-07) | - | -3.11<br>(1.51E-12) | MFP1 paralog. Potential role in slowing MSP polymerization |
| *msd-3* | -2.14<br>(4.38E-04) | - | -2.73<br>(1.51E-12) | MFP1 paralog. Potential role in slowing MSP polymerization |
| *smz-1* | - | - | -3.48<br>(6.40E-08) | Chromatin bound protein. RNAi causes meiotic metaphase arrest. |
| *smz-2* | -5.00<br>(1.43E-30) | - | -5.16<br>(6.59E-33) | Chromatin bound protein. RNAi causes meiotic metaphase arrest. |
| *elt-1* | - | -1.12<br>(4.77E-02) | -1.21<br>(3.78E-02) | Transcription factor that regulates *msp* gene expression. Regulated by *spe-44* in Kulkarni et al., (2012) microarray. |
| *msp-49* | - | -1.51<br>(9.29E-03) | -2.57<br>(2.55E-08) | Major sperm protein gene. Regulated by *spe-44* in Kulkarni et al., (2012) microarray. |
| *msp-63* | - | -2.17<br>(8.50E-06) | -1.56<br>(7.07E-03) | Major sperm protein gene. Regulated by *spe-44* in Kulkarni et al., (2012) microarray. |
| *spe-4* | - | - | -1.74<br>(2.64E-02) | Required for cytoplasmic partitioning. |
| *spe-9* | - | -3.74<br>(2.23E-04) | -5.71<br>(2.96E-07) | Membrane protein required for fertilization. |
| *spe-11* | - | - | -1.23<br>(1.31E-02) | Prevents polyspermy and required for eggshell formation |
| *spe-15* | - | - | -2.61<br>(1.81E-07) | Myosin IV homolog required for cytoplasmic partitioning. |
| *spe-18* | - | -1.59<br>(5.97E-03) | -5.50<br>(8.95E-35) | Promotes MSP assembly within fibrous bodies. Regulated by *spe-44* in Kulkarni et al., (2012) microarray. |
| *spe-42* | - | -2.04<br>(3.31E-02) | -3.35<br>(3.13E-05) | Membrane protein required for fertilization. |
| *spe-43* | - | -3.78<br>(3.67E-04) | -6.99<br>(1.80E-05) | Required for sperm activation. |
| *spe-45* | - | -4.44<br>(1.13E-06) | -4.77<br>(1.56E-07) | Required for spermatozoa fusion with the oocyte plasma membrane. |
| *spe-49* | - | -3.25<br>(7.53E-05) | -6.25<br>(3.48E-09) | Membrane protein required for fertilization. |
| *nsph-1.1* | - | - | -2.18<br>(2.90E-03) | MFP1 paralog. Potential role in promoting MSP polymerization |
| *nsph-2* | - | -1.70<br>(1.05E-03) | - | MFP1 paralog. Potential role in promoting MSP polymerization |
| *nsph-3.1* | - | -1.26<br>(3.06E-02) | -1.53<br>(3.59E-03) | MFP1 paralog. Potential role in promoting MSP polymerization |
| *nsph-3.2* | -1.52<br>(1.50E-02) | -2.26<br>(4.17E-07) | -4.94<br>(1.59E-35) | MFP1 paralog. Potential role in promoting MSP polymerization |
| *nsph-4.1* | -8.32<br>(3.71E-67) | -5.97<br>(1.16E-40) | -8.35<br>(9.11E-68) | MFP1 paralog. Potential role in promoting MSP polymerization |
| *nsph-4.2* | -8.55<br>(1.00E-72) | -6.11<br>(6.61E-45) | -8.57<br>(2.36E-73) | MFP1 paralog. Potential role in promoting MSP polymerization |

There were 326 differentially-regulated genes unique to NHR-23+SPE-44-depleted males, consistent with the additive phenotype when we deplete both SPE-44 and NHR-23 (Ragle *et al*. 2020)(Fig. 2). These genes included 14 *msp* genes, as well as *smz-1,* the paralog of NHR-23-regulated *smz-2* (Table 1, Table S7). There were also three *spe* genes downregulated when both NHR-23 and SPE-44 were depleted (Table 1). The presenilin paralog SPE-4 localizes to FB-MOs and is required for cytoplasmic partitioning during spermatogenesis (L’Hernault *et al*. 1988; Arduengo *et al*. 1998). A *spe-4* promoter reporter was used by Kulkarni *et al*. to validate their microarray data and confirm that it was not regulated by *spe-44.* The paternal effect factor SPE-11 prevents polyspermy and is required for eggshell formation (Hill *et al*. 1989), and SPE-15, a myosin VI homolog expressed in primary spermatocytes, is required for asymmetric partitioning of components such as FB-MOs and mitochondria during residual body formation (Kelleher *et al*. 2000; Hu *et al*. 2019).

Fifteen differentially-regulated genes were common to NHR-23-, SPE-44-, and NHR-23+SPE-44-depleted animals (Fig. 2). Three of these genes were MFP2 paralogs (*nsph-3.2, nsph-4.1, nsph-4.2*) (Table 1). MFP2 enhances MSP polymerization in *Ascaris* sperm, countering the activity of MFP1 (Grant *et al*. 2005), and there are eight MFP2 paralog genes in *C. elegans* which have been assigned the nematode-specific peptide family, group H (*nsph*) gene class (Fig. S2)(Rödelsperger *et al*. 2021). *nsph-2* is downregulated when SPE-44 is depleted, *nsph-3.1* is found in both the SPE-44 and NHR-23+SPE-44 depleted differentially-regulated genes, and *nsph-1.1* is down-regulated only when both NHR-23 and SPE-44 are depleted (Table 1). Two *nsph* genes (*1.2, 4.3*) are not found in any of the datasets. We note that *nsph-4.3* has a large deletion that removes 500 bp of sequence, though a predicted protein product can still be produced (Fig S2, S3). There is also a pseudogene MFP2 paralog (ZK546.7) down-regulated following NHR-23-, SPE-44, and NHR-23+SPE-44 depletion. This pseudogene is related to *nsph-4.1* and *nsph-4.2* but an insertion causes a frameshift. An initiation codon further downstream could produce a valid coding sequence, so this pseudogene may be worth future exploration (Fig. S3).

### NHR-23 and SPE-44 targets are enriched in phosphatase genes

To gain insight into how NHR-23 and SPE-44 promote spermatogenesis, we performed gene ontology (GO) analysis on the differentially-regulated genes to identify enriched biological processes. Phosphatases were enriched in all three depletion conditions (Table 2, S5-7). The NHR-23-regulated genes were also enriched in genes involved in ATP binding, though these genes overlapped with the phosphatase genes and cytoskeletal protein binding (Table S8). SPE-44-regulated genes were enriched in genes involved in DNA replication (origin binding, helicase activity, single-strand DNA-dependent ATPase), carbohydrate-binding, glutamate-ammonia ligase activity, and calcium ion-binding (Table S9). In addition to being enriched in phosphatase-related GO terms, the NHR-23+SPE- 44-regulated genes were also enriched in kinase genes, NADPH reductase activity, FFAT motif binding, drug binding and transaminase activity (Table S10). Together, these data implicate NHR-23 and SPE-44 in regulating signal transduction by kinases and phosphatases, which has been shown to be an important regulatory mode following transcriptional quiescence in spermatocytes (Muhlrad and Ward 2002; Bae *et al*. 2009; Wu *et al*. 2012).

**Table 2.** NHR-23-regulated kinases and phosphatases.

| Gene | Log2 fold-change | Class | Ortholog |
| --- | --- | --- | --- |
| *ZK596.2* | -9.01 | Predicted serine/threonine kinase | Tau-tubulin kinase 2 (human); Casein kinase I delta ortholog |
| *W03G9.5* | -8.19 | Predicted serine/threonine kinase | Casein kinase delta (human) and Casein kinase I (yeast) |
| *F47B3.2* | -4.17 | Predicted tyrosine phosphatase | Tyrosine-protein phosphatase non-receptor type 6 isoform 3 (mammals) |
| *F55H12.5* | -4.04 | Predicted tyrosine phosphatase | Receptor-type tyrosine-protein phosphatase T (mammals) |
| *F47B3.7* | -4 | Predicted tyrosine phosphatase | Tyrosine-protein phosphatase non-receptor type 6 isoform 3 (mammals) |
| *ZK757.2* | -3.49 | Predicted tyrosine/serine/threonine phosphatase | Dual specificity phosphatase 14 (human) |
| *D2045.7* | -3.05 | Predicted serine/threonine kinase | Serine/threonine kinase 16 (human) |
| *F52H3.6* | -2.7 | Predicted serine/threonine phosphatase | Isoform Gamma-1 of Serine/threonine-protein phosphatase PP1-gamma catalytic subunit (humans) |
| *ttbk-5* | -2.01 | Predicted serine/threonine kinase | Tau-tubulin kinase 2 (human); Casein kinase I delta ortholog |

### NHR-23 regulates uncharacterized kinases and MFP2 paralogs

We focused on the NHR-23-regulated genes as they were few in number so therefore amenable to a candidate-based knock-out approach. In addition to the aforementioned phosphatase and *nsph* genes, NHR-23 also regulated four uncharacterized kinases (Table 2). Three of these kinases were predicted to be Casein Kinase I orthologs. W03G9.5 and C04G2.2 belong to the Tau Tubulin Kinase-Like subgroup, while ZK596.2 belongs to the nematode *spe-6* subgroup (Manning 2005). We engineered predicted null mutant alleles through insertion of a “STOP-IN” cassette containing premature termination codons in all three reading frames in the first or second exon of targeted genes (Wang *et al*. 2018). We generated mutants in one phosphatase (*ZK757.2*), two kinases (*ttbk-5, ZK596.2*), and three MFP2 paralogs (*nsph-2, nsph-3.2, nsph-4.3*) all of which were viable with wild-type brood sizes (Table 3). *nsph-4.1* and *nsph-4.2* are predicted to produce identical proteins and the genes only differ by a single nucleotide in an intron and have 1231 bases of identical upstream promoter sequence. This sequence similarity made genotyping a STOP-IN insertion challenging, so we deleted the entire gene for each using crRNAs that bound outside the conserved sequence for each gene. Each mutant had a wild-type brood size and no obvious increase in unfertilized oocytes (Table 3). These data could suggest that these genes are dispensable for fertility, redundant with other genes, or the STOP-IN cassette is not creating a null. We created *nsph-2; nsph-1.1* and *nsph-4.1, nsph-4.2* double mutants, both of which had wild-type brood sizes. These data could suggest that the *nsph* paralogs are dispensable for *C. elegans* spermatogenesis or that extensive redundancy exists.

**Table 3.** Brood size data.

| Genotype | Allele class | Predicted gene function | Brood size (±sd) | % unfertilized eggs |
| --- | --- | --- | --- | --- |
| N2 (wt) | N/A | N/A | 230±91 | 3.2 (n=2891) |
| *nsph-2(wrd57)* | PTC | MFP2 paralog | 301±43 | 2.75 (n=2184) |
| *nsph-3.2(wrd56)* | PTC | MFP2 paralog | 277±39 | 2.34 (n=3420) |
| *nsph-2(wrd57); nsph-3.2(wrd56)* | PTC | MFP2 paralog | 293±39 | 1.5 (n=3276) |
| *nsph-2(wrd57); nsph-1.1(wrd67)* | PTC | MFP2 paralog | 185±40 | 1.07 (n=2241) |
| *nsph-4.3(wrd74)* | PTC | MFP2 paralog | 277±49 | 3.08 (n=3445) |
| *nsph-4.1(wrd92)* | Whole gene deletion | MFP2 paralog | 228±45 | 5.75 (n=2677) |
| *nsph-4.2(wrd93)* | Whole gene deletion | MFP2 paralog | 219±35 | 4.15 (n=2750) |
| *nsph-4.1(wrd92); nsph-4.2(wrd97)* | Whole gene deletion | MFP2 paralog | 260±49 | 4.64 (n=3296) |
| *ttbk-5(wrd72)* | PTC | Kinase | 259±25 | 6.84 (n=3373) |
| *ZK596.2(wrd71)* | PTC | Kinase | 231±80 | 2.93 (n=2931) |
| *ZK757.2(wrd77)* | PTC | Phosphatase | 242±17 | 3.31 (n=2999) |

### NHR-23-depletion causes NSPH-2 and MSD-1 localization defects

As previously described, both NSPH-2 and MSD-1 antibodies label fibrous bodies (FBs) and co-localize with MSP antibody labeling in wildtype 1° spermatocytes ((Morrison *et al*. 2021), Fig. 3A,B). However in previous studies (Kulkarni *et al*. 2012; Ragle *et al*. 2020), we found that single and double NHR-23 and SPE-44 germline depletion have distinct effects on FB-MO formation. When labeled with the MO differentiation marker 1CB4, NHR-23 depleted spermatocytes exhibit distinct labeled structures, whereas 1CB4 labeling is diffuse in SPE-44 depleted spermatocytes and absent in the NHR-23+SPE-44 depleted spermatocytes (Ragle *et al*. 2020). In all cases, MSP fails to assemble into FBs in spermatocytes (Kulkarni *et al*. 2012; Ragle *et al*. 2020). Therefore, we wondered how depleting NHR-23, SPE-44, and NHR-23+SPE-44 from the germline would affect the localization of other FB components such as NSPH-2 and MSD-1. In each of our depletion strains, germline labeling with anti-NSPH-2 and anti-MSD-1 was dim and diffuse, compared to the bright puncta labeled in control germlines, indicating that assembly of NSPH-2 and MSD-1 into FBs was impaired in the absence of NHR-23, SPE-44, or both. Although our RNA-seq data suggests that SPE-44 and not NHR-23 regulates *nsph-2* (Table 1), terminal SPE-44 depleted spermatocytes exhibited NSPH-2 aggregates that were not observed in either NHR-23 or the double depleted strains (Fig. 3A). Due to this unexpected phenotype, we also investigated *spe-44* null (*ok1400*) mutants (Kulkarni et al., 2012). Auxin-inducible depletion of target proteins is robust, but may not be fully penetrant in some cases, so the null mutant provides additional insight into *spe-44*’s role in FB assembly. Consistent with our RNA-seq results, anti-NSPH did not label germlines in *spe-44(ok1400)* animals (Fig. 3A’). As predicted from our RNA-seq data (Table 1), MSD-1 levels were most diminished in NHR-23 and NHR-23+SPE-44 depleted spermatocytes, whereas terminal SPE-44 depleted spermatocytes exhibited dim but detectable MSD-1 punctae (Fig. 3B). In both NHR-23 and SPE-44 knockdowns, the MSD-1 antibody labeled centrosomes (arrows), but the significance of this pattern is unclear (Fig. 3B). This data indicates that NHR-23 is either required for localization of NSPH-2 and MSD-1 to the MOs or for the stable expression of NSPH-2 and MSD-1, which is consistent with our previous finding that NHR-23 depletion prevents MSP loading into FBs (Ragle *et al*. 2020).

**Fig. 3.**
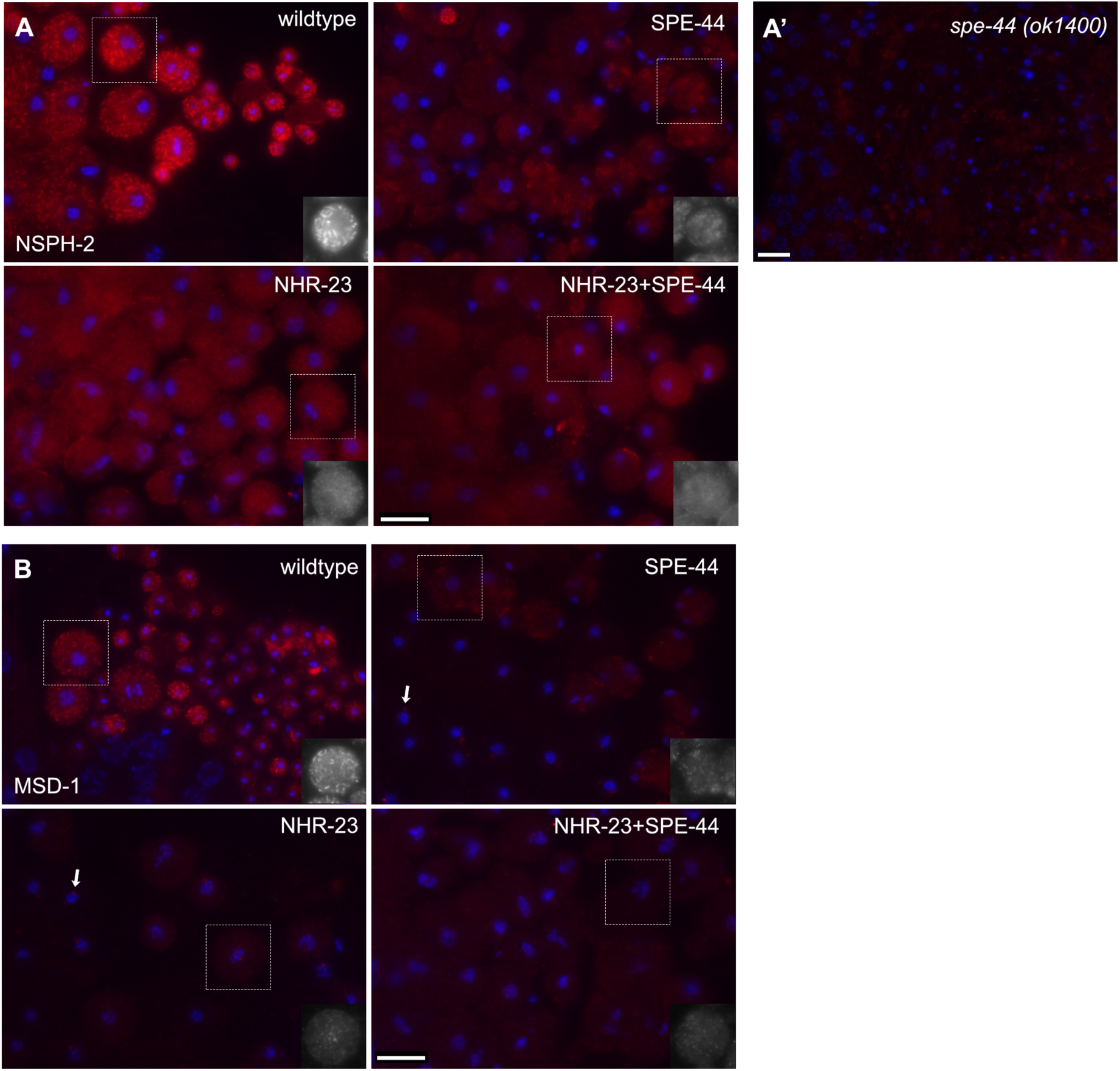
NSPH-2 and MSD-1 in NHR-23, SPE-44, and NHR-23+SPE-44 depleted spermatocytes. Monolayer of isolated spermatocytes; and, in wildtype, budding figures and spermatids labeled with DAPI (blue) and either anti-NSPH-2 antibody (A) or anti-MSD-1 antibody (B) (red). Images are oriented with earlier stages on the left. In the images shown, wildtype spermatocytes are from JDW92 males that contain the *nhr-23* degron constructs but were not auxin-treated (A,B). The other images are from males in which the indicated transcription factor has been depleted by auxin-mediated degradation. Images are representative of multiple preparations, but images shown are from the same preparation and taken with the same exposures without additional adjustments. (A’) Monolayer of isolated spermatocytes from *spe-44(ok1400)* null mutant labeled with anti-NSPH-2 antibody. (B) Arrows show potentially non-specifically labeled centrosomes. In A and B, black and white inserts of indicated spermatocytes are shown for clarity. Scale bars = 10 microns.

### SPE-6 patterns are differentially affected in single and double knockdowns

Given our finding that NHR-23 regulates three uncharacterized Casein Kinase I orthologs, W03G9.5, C04G2.2, and ZK596.2, which belong to the nematode *spe-6* subgroup; we decided to investigate the localization of SPE-6, the better characterized casein kinase 1 ortholog, in our depletion strains. In *C. elegans*, *spe-6* encodes one of a large family of casein kinase 1 proteins (Muhlrad and Ward 2002). Like NHR-23 and SPE-44 depletion, SPE-6 null mutations result in primary spermatocyte arrest (Varkey *et al*. 1993). During wildtype spermatogenesis, SPE-6 shifts from a diffuse plus particulate pattern in spermatocytes to being localized around the condensed haploid chromatin mass during the budding division that follows anaphase II (Peterson *et al*. 2021; Fig. 4A). Although *spe-6* transcript levels were not significantly altered in either the single or double depletion strains (Table S1), we tested whether either the SPE-6 localization patterns or levels of abundance would be differentially altered when we used anti-SPE-6 antibody to label the proximal germlines of affected males. In NHR-23 depleted spermatocytes, SPE-6 remains in a diffuse plus particulate pattern, consistent with its early meiosis I arrest (Fig. 4B). In contrast, in SPE-44 depleted spermatocytes, SPE-6 clumps around the chromatin masses in a post-meiotic-like pattern even though the spermatocytes never physically divide to form haploid spermatids (Fig. 4C, Kulkarni *et al*. 2012). Anti-SPE-6 labeling in NHR-23+SPE-44 depleted germlines exhibited two distinct phenotypes: some were indistinguishable from single depletion while others failed to label with SPE-6 antibodies (Fig. 4D). These data suggest that the developmental program of restructuring the sperm chromatin continues upon SPE-44 depletion but not upon NHR-23 depletion and that SPE-6 expression is lost upon NHR-23+SPE-44-depletion.

**Fig. 4.**
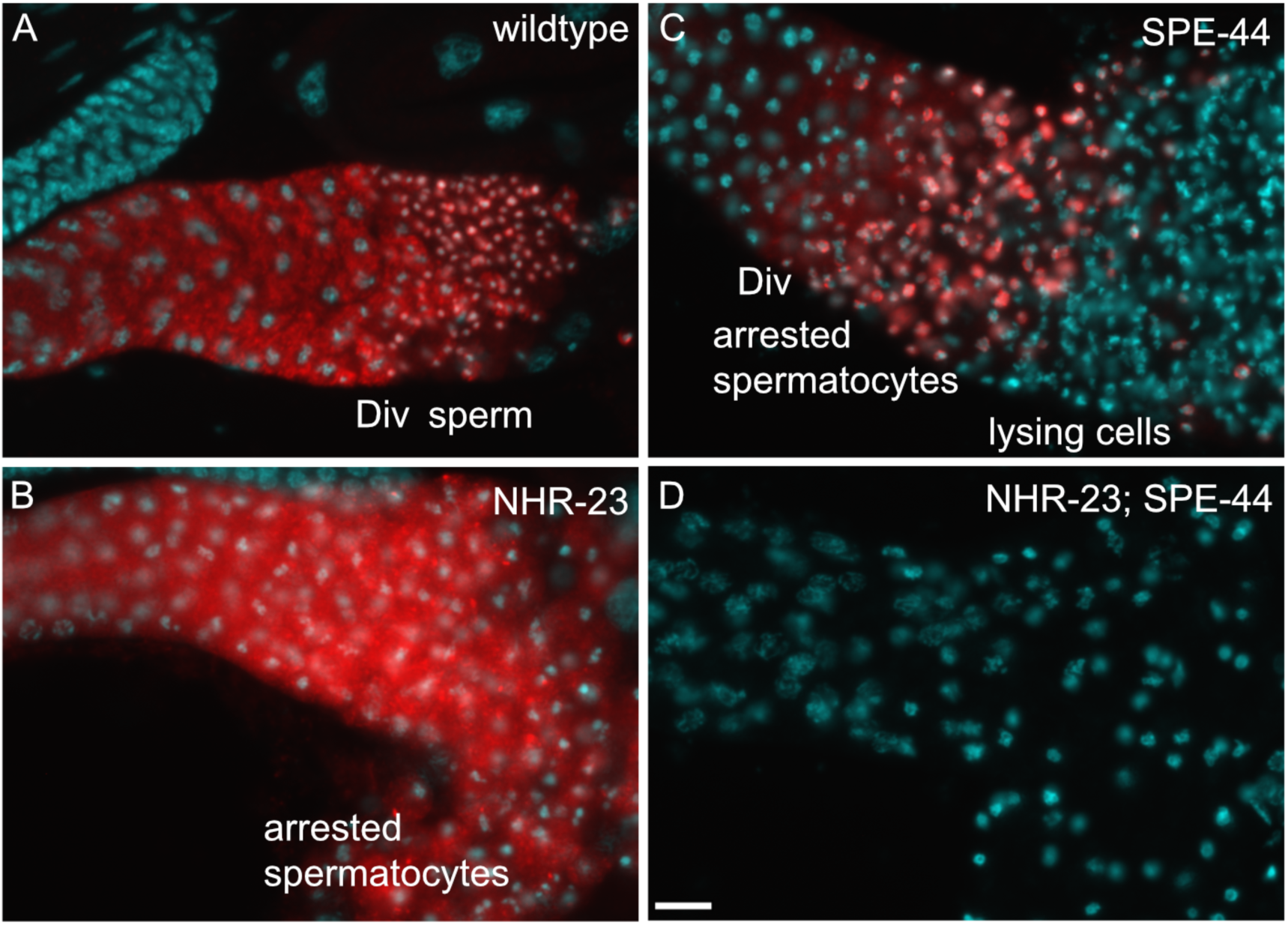
SPE-6 patterns in NHR-23, SPE-44, and NHR-23+SPE-44 depleted spermatocytes. Proximal regions of male gonads labeled with DAPI (cyan) and anti-SPE-6 antibody (red). Images are oriented with earlier stages on the left. (A) Wildtype are auxin-treated *him-5* males. (B-C) The other images are from males in which the indicated transcription factor has been depleted by auxin-mediated degradation. Regions of the gonad are labeled including the meiotic division zone (DIV). Images are representative of multiple preparations, but images shown are from the same preparation and taken with the same exposures without additional adjustments. Scale bars = 10 microns.

## DISCUSSION

This work identifies genes differentially regulated following NHR-23-depletion in male germlines, providing insight into how NHR-23 promotes spermatogenesis. We also identify genes specifically differentially expressed when both NHR-23 and SPE-44 are depleted in male germlines, providing an entry point to understanding the synthetic phenotype when these two transcription factors are depleted (Ragle *et al*. 2020). NHR-23 depletion causes defective localization of NSPH-2 and MSD-1 to FBs, implicating NHR-23 in regulating MSP polymerization dynamics. Although NHR-23 and SPE-44 don’t appear to transcriptionally regulate the Casein kinase *spe-6,* double depletion of these transcription factors causes a loss of SPE-6 antibody labeling, suggesting that SPE-6 protein levels may be compromised when NHR-23 and SPE-44 are absent from the germline.

### NHR-23 regulates few genes with known roles in spermatogenesis

Relatively few genes were differentially regulated following NHR-23-depletion (Table S5; 70 genes) compared to SPE-44-depletion (Table S6; 811 genes), NHR-23+SPE-44- depletion (Table S7; 554 genes), and *nhr-23(RNAi)* in L2 larvae (Kouns *et al*. 2011). Aside from *smz-2*, none of the NHR-23-regulated genes have been experimentally confirmed to play roles in spermatogenesis.

Terminally arrested spermatocytes from NHR-23 depleted animals arrest early in meiosis and contain distended MOs, similar to *spe-6(lf)* and *wee-1.3(gf)* mutants (Varkey *et al*. 1993; Lamitina and L’Hernault 2002), yet NHR-23 does not regulate these genes (Table S1, S5). The dearth of known spermatogenesis regulators is consistent with NHR-23 regulating a novel pathway promoting sperm development. As NHR-23 regulates several uncharacterized kinases and phosphatases, one possibility is that NHR-23 controls meiotic progression and FB-MO biogenesis through kinase-mediated signal transduction.

The orthologs of two of the kinases identified in our RNA-seq dataset (*ttbk-5, ZK596.2*) are implicated in mammalian microtubule dynamics and yeast vesicular traffic, meiotic exit, and kinetochore attachment in meiosis II (Petronczki *et al*. 2006; Goetz *et al*. 2012; Lee *et al*. 2017). While our STOP-IN insertions didn’t have phenotypes, there could be redundancy or genetic compensation, a point which we will expand on in the next section.

### NHR-23 is necessary for MSD-1 and NSPH-2 localization

Our immunostaining revealed that NHR-23-depletion caused a loss of MSD-1 and NSPH-2 localization in spermatocytes. The orthologs of these factors (MFP1 and MFP2) in *Ascaris* are specifically known to control MSP polymerization within the pseudopod of crawling sperm (Buttery *et al*. 2003; Grant *et al*. 2005) but since they are also present in FBs (Morrison et al., 2021), their downregulation upon NHR-23 depletion might account for the observed defect in MSP loading into FB-MOs (Ragle *et al*. 2020). The effect of NHR-23 on NSPH-2 localization was surprising as this gene was not differentially regulated following NHR-23 depletion (Tables 1, S5). *nsph-2* was downregulated following SPE-44 depletion (Tables 1, S5); and by immunocytology, the protein expression appeared slightly reduced in SPE-44-depleted germlines and significantly reduced in *spe-44(ok1400*) null mutants (Fig. 3). This data suggests that NHR-23 could promote NSPH localization to FB-MOs. There are four uncharacterized MSD1 paralogs in *C. elegans*, three of which are regulated by NHR-23 (Tables 1, S5). There are eight MFP2 paralogs, which are also uncharacterized. Three of these paralogs are regulated by NHR-23 while five are SPE-44-regulated (Table 1). No phenotype has been previously reported for the *C. elegans* MFP1 or MFP2 paralogs, though redundancy may be responsible since these paralogs appear to have arisen through duplication. We have inactivated six of the MFP2 paralogs either singly or in double mutant combinations and have seen no difference in fertility. Three possible explanations for the lack of phenotype are: i) that these genes do not play a role in *C. elegans* spermatogenesis; ii) extensive genetic redundancy exists and we will have to inactivate more paralogs, if not all, to produce a phenotype; or iii) that our knock-out approach caused genetic compensation. We favor the second and third possibilities. Our initial approach to inactivating candidate NHR-23- and SPE-44-regulated genes was to insert premature termination codons in all three reading frames, as this approach was reported to produce null phenotypes and requires a single crRNA/sgRNA and repair oligo (Wang *et al*. 2018). However, given the homology between the MFP2 paralogs, they may be subject to genetic compensation. This phenomenon, reported in zebrafish, mammalian cells, and *C. elegans*, results in upregulation of paralogous genes when an mRNA with a premature termination codon is translated in the cytoplasm (Rossi *et al*. 2015; El-Brolosy *et al*. 2019, 2019). Mutations that prevent mRNA production, such as promoter deletions or whole gene deletions do not trigger this response (El-Brolosy *et al*. 2019), so our *nsph-4.1, nsph-4.2* single and double mutants should not be subject to this phenomenon but the other knock-outs could be. Going forward, it will be important to either systematically delete MFP1 and MFP2 orthologs or use a multiplexed piRNA-based knockdown approach to determine whether these genes play roles in *C. elegans* MSP polymerization dynamics (Priyadarshini *et al*. 2022).

### A high-confidence set of *spe-44-*regulated genes

There was a surprising lack of overlap between the SPE-44-regulated genes in our RNA-seq dataset and differentially regulated genes in a *spe-44* mutant microarray dataset (Kulkarni *et al*. 2012), though the genes common to both datasets are high-confidence *spe-44* targets. Our data was generated from adult males, while the *spe-44* mutant dataset was generated from L4 males, so it is possible that differences in developmental stage impacted the results. While RNA-seq is reported to have a wider quantitative range in detecting changes in expression levels, a comparison of microarrays vs RNA-seq on rat cDNA reported a 78% overlap in differentially regulated genes (Rao *et al*. 2019). The difference could be biological as the *spe-44* mutant is a predicted null, while auxin-mediated protein depletion frequently produces hypomorphic phenotypes as we also observed in our NSPH-2 immunocytology. The genes common to both datasets include *elt-1*, a transcription factor that acts downstream of SPE-44 to regulate MSP gene expression, and *ceh-48* (del Castillo-Olivares *et al*. 2009; Kulkarni *et al*. 2012). *ceh-48* is an uncharacterized ONECUT homeobox transcription factor ortholog that Kulkarni *et al*. (2012) identified as sperm-enriched. A deletion mutant is reported to be fertile, but displays male tail defects (Kim *et al*. 2016). *spe-18, which* promotes MSP assembly within FBs, and the MFP2 paralog *nsph-2* were also common to both datasets (Price *et al*. 2021). Exploring the high confidence target will provide insight into how SPE-44 promotes FB-MO assembly and meiosis II.

### SPE-6 labeling is lost in NHR-23+SPE-44 depleted germlines

SPE-6 is a casein kinase that plays broad roles in spermatogenesis including FB assembly, meiotic progression, and repression of precocious sperm activation (Varkey *et al*. 1993; Muhlrad and Ward 2002; Price *et al*. 2021; Peterson *et al*. 2021). *spe-6* was not differentially-regulated in any of our datasets, despite the overlap of the NHR-23-depletion and *spe-6(lf)* meiotic arrest and distended FB-MO phenotype (Varkey *et al*. 1993; Ragle *et al*. 2020). SPE-6 labeling in NHR-23-depleted animals appeared in a punctate pattern, as well as diffuse in the cytosol, as expected in metaphase I arrested spermatocytes (Ragle *et al*. 2020; Fig 4C). In SPE-44-depleted germlines, SPE-6 labeling appeared similar to a post-meiotic pattern, despite the meiosis II arrest and lack of spermatocytes (Kulkarni *et al*. 2012; Fig. 4B). These data suggest that progression through metaphase I is required for SPE-6 to localize around chromatin after meiosis, and that this localization and presumably SPE-6 pathway is insensitive to SPE-44 depletion and the meiosis II arrest. Surprisingly, NHR-23+SPE-44 double-depletion caused a complete loss of SPE- 6 labeling, another example of a synthetic phenotype caused by depletion of these transcription factors (Fig. 4D). *spe-6* transcript levels were not significantly affected by NHR-23+SPE-44 depletion (Table S1), indicating that this SPE-6 down-regulation is not transcriptional. Possible mechanisms could include impaired translation of *spe-6* mRNA or an increase in SPE-6 turnover.

### Future perspectives

We previously discovered a novel role for NHR-23 in spermatogenesis and a synthetic interaction between NHR-23 and SPE-44 (Ragle *et al*. 2020). We also proposed that a yet-undetermined pathway promoted spermatogenesis independently of NHR-23 and SPE-44 (Ragle *et al*. 2020). In agreement with this assertion, there are a large number of sperm-enriched genes not found in our differentially-regulated gene lists, and there are genes only differentially regulated following NHR-23+SPE-44 depletion, consistent with synthetic interaction. Numerous experimentally validated *spe* genes are not regulated by NHR-23 or SPE-44. Our RNA-seq data provides an entry point to understanding transcriptional control of *C. elegans* spermatogenesis and how NHR-23 and SPE-44 interact to promote FB-MO biogenesis.

## Competing interests

The authors declare no competing or financial interests.

## Acknowledgements

This work was funded by grants from the National Science Foundation (NSF) Division of Molecular and Cellular Biosciences (CAREER award 1942922) to J.D.W. and National Institutes of Health (R15GM-096309) to D.C.S. Some strains were provided by the Caenorhabditis Genetics Center, which is funded by the NIH Office of Research Infrastructure Programs [P40 OD010440].

---

**Fig. S1.**
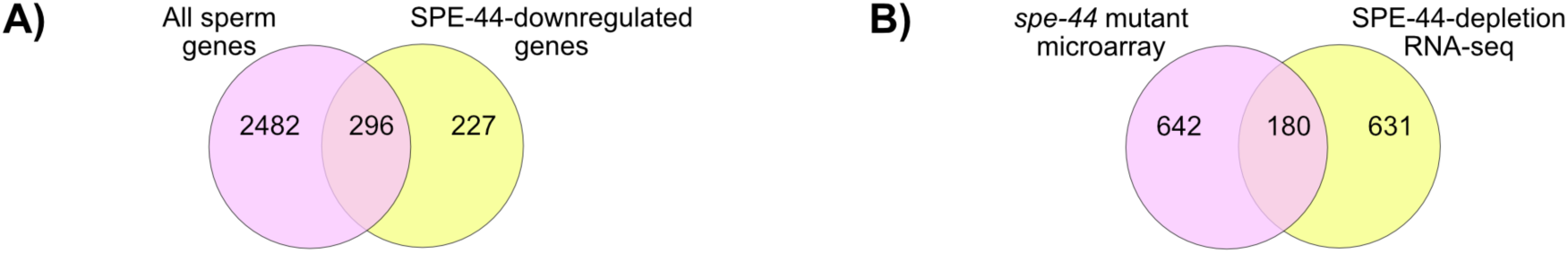
Overlaps between SPE-44-downregulated genes and sperm-enriched genes, and comparison of SPE-44 RNA-seq and *spe-44* microarray datasets. (A) Venn diagram of down-regulated genes following SPE-44 depletion and sperm-enriched genes. (B)Venn diagram of genes differentially regulated following SPE-44 depletion (this study) and a previous microarray experiment comparing gene expression in L4 males between spe-44 null mutants and wild-type animals (Kulkarni et al. 2012).

**Figure S2.**
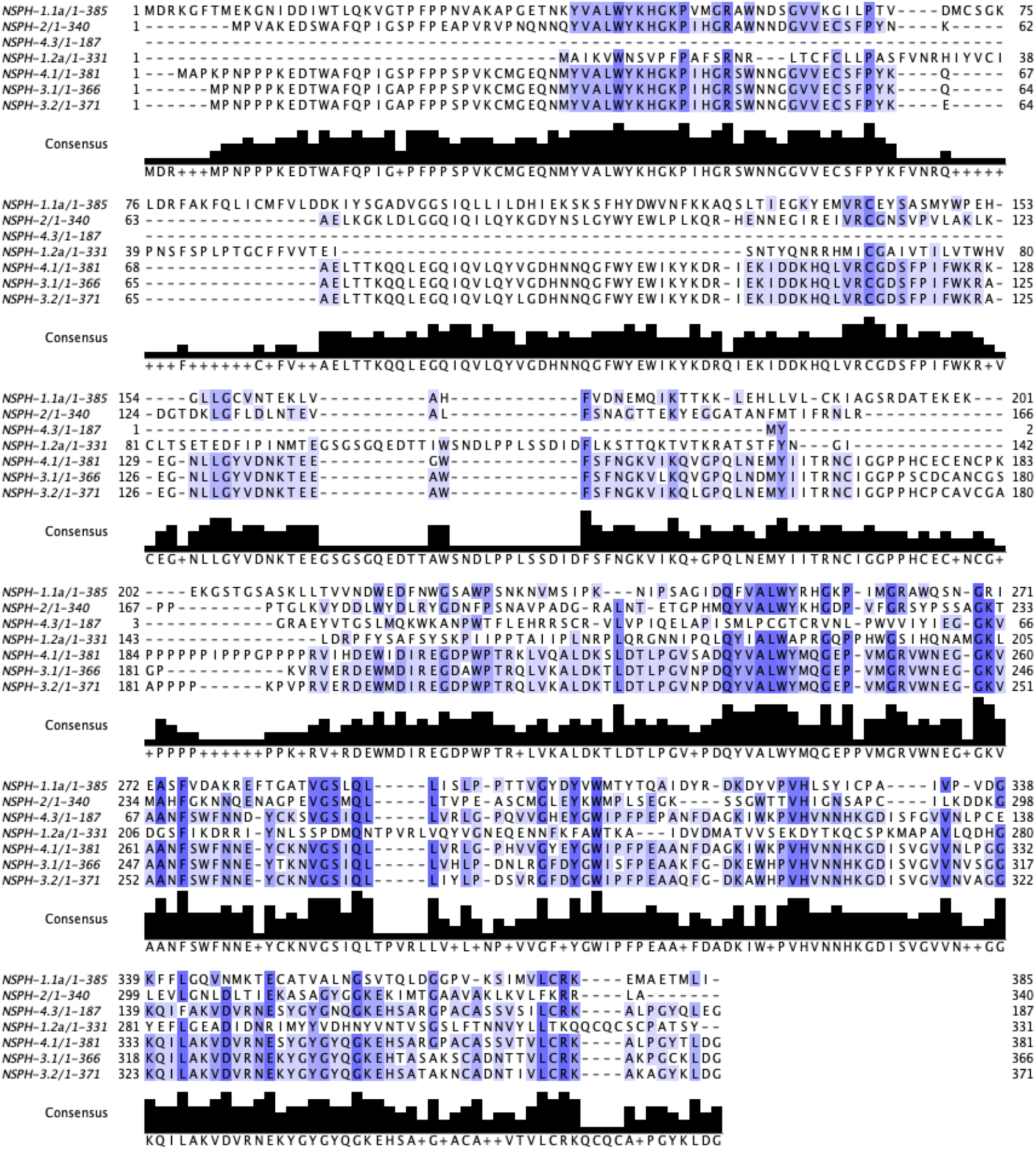
NSPH paralog alignment. An alignment produced with Clustal Omega (Goujon et al., 2010; Sievers et al., 2011) and visualized with JalView2 (Waterhouse et al., 2009). Protein identity for each row and total length in amino acids is provided in the left column. To the left and right of the alignment are amino acid position references. The consensus sequence is shown beneath each row. Only the sequence for NSPH-4.1 is shown as NSPH-4.2 has identical sequence and its protein product is annotated as NSPH-4.1 in Wormbase.

**Figure S3.**
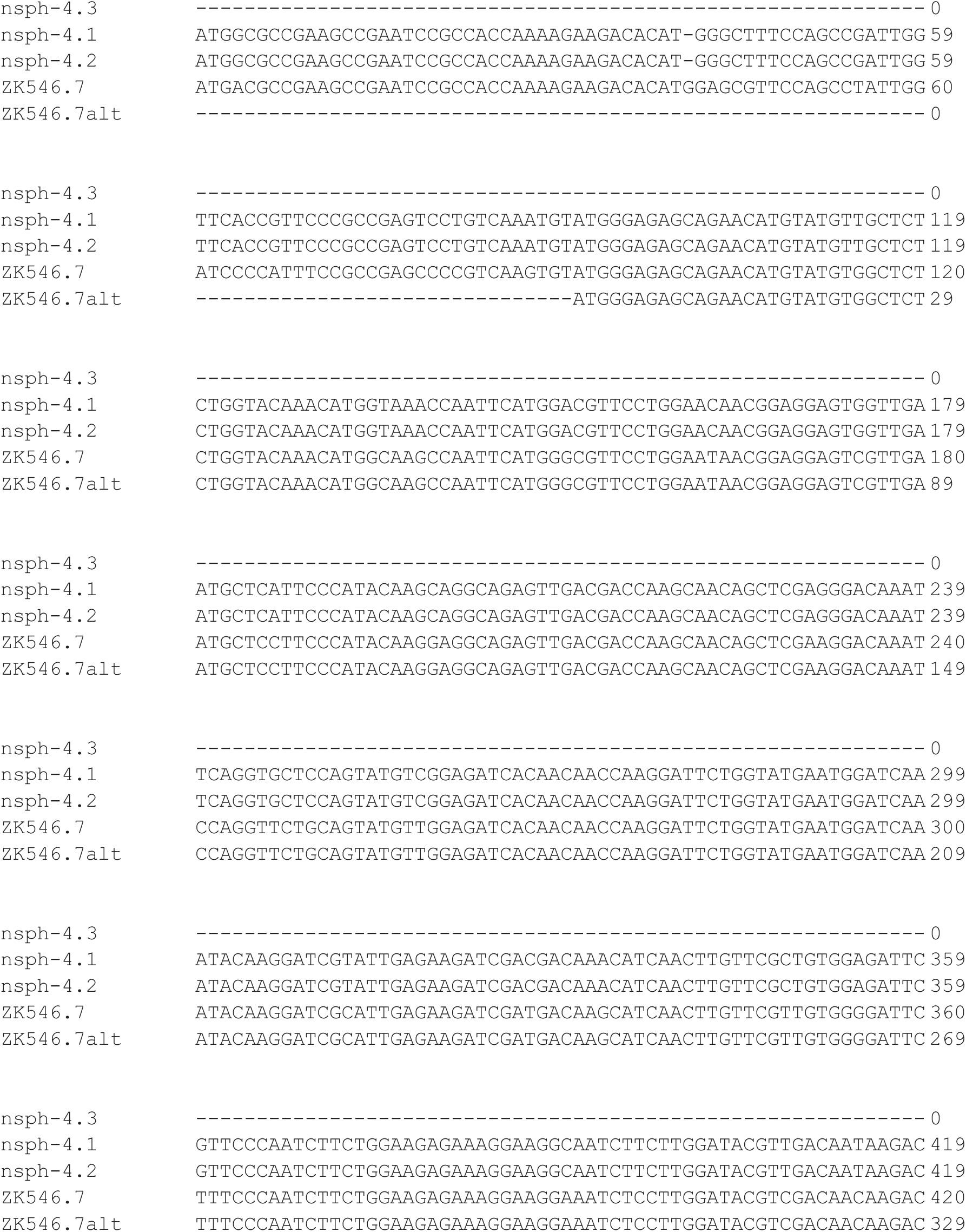
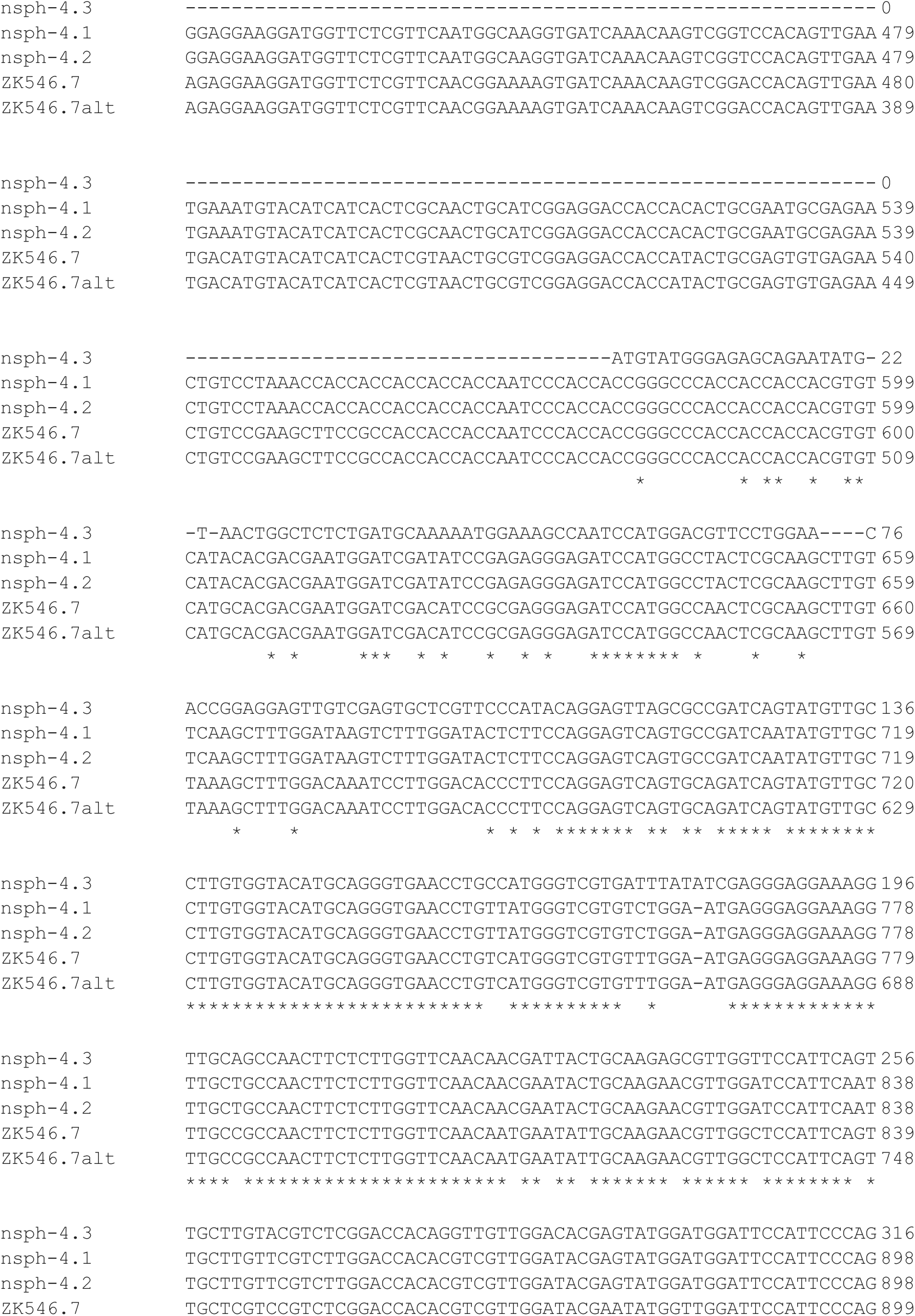
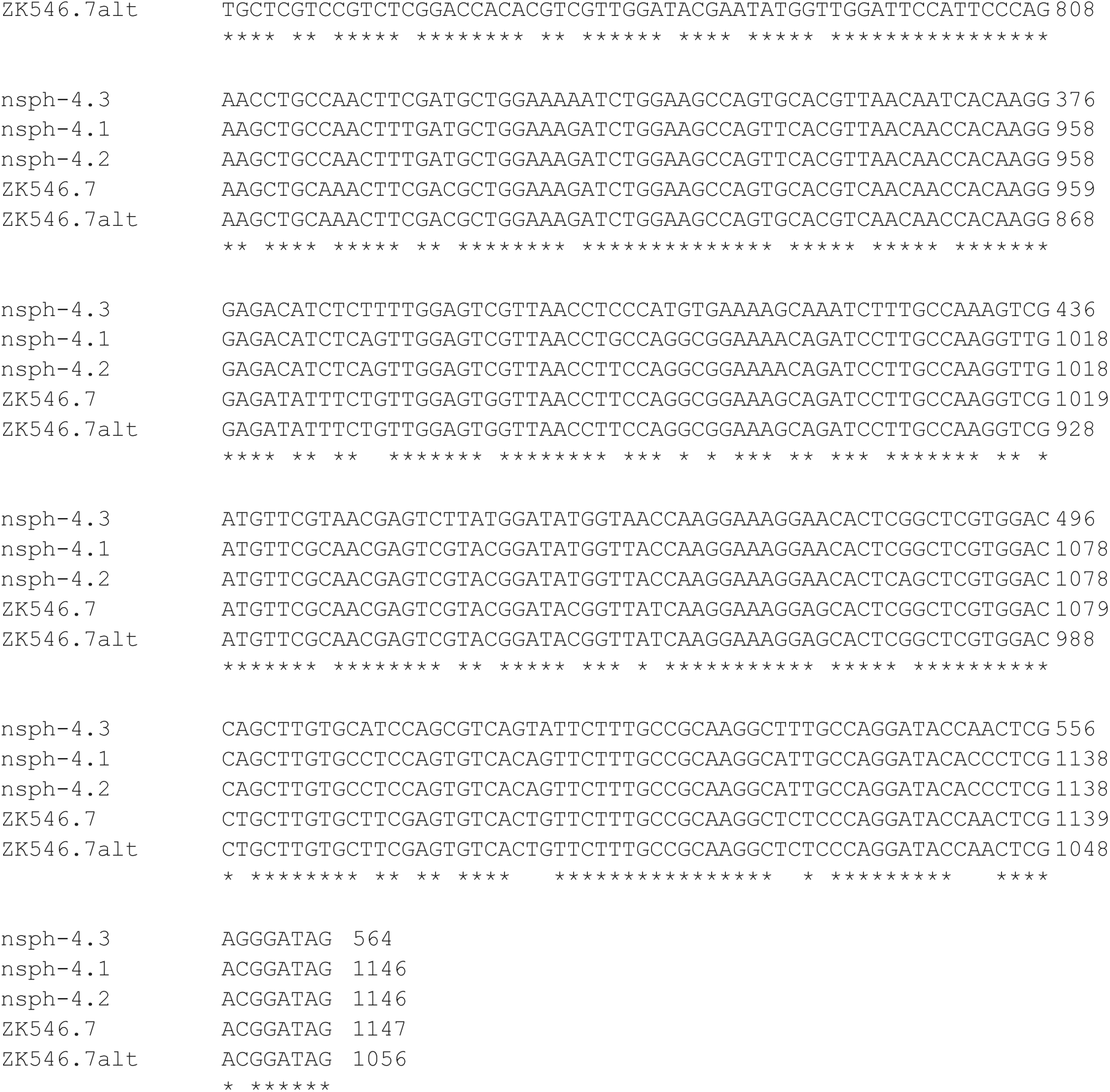
MFP2 genomic DNA alignment. An alignment produced with Clustal Omega (Goujon et al., 2010; Sievers et al., 2011). cDNA identity for each row and total length in basepairs is provided in the right column. An alternate potential *ZK596.2* reading frame starting 92 bp downstream from the conserved start codon is shown in *ZK596.2alt*.

